# CAMIO for deletion analysis of endogenous DNA sequences in multicellular organisms

**DOI:** 10.1101/658088

**Authors:** Hui-Min Chen, Jorge Garcia Marques, Ken Sugino, Dingjun Wei, Rosa Linda Miyares, Tzumin Lee

## Abstract

The genome is the blueprint for an organism. Interrogating the genome, especially locating critical cis-regulatory elements, requires deletion analysis. This is conventionally performed using synthetic constructs, making it cumbersome and non-physiological. Thus, we created <u>C</u>as9-mediated <u>A</u>rrayed <u>M</u>utagenesis of Individual <u>O</u>ffspring (CAMIO) to achieve high-throughput analysis of native DNA. CAMIO utilizes CRISPR that is spatially restricted to generate independent deletions. Controlled by recombination, a single guide RNA is stochastically chosen from a set targeting a specific DNA region. Combining two sets increases variability, leading to either indels at 1-2 target sites or inter-target deletions. Cas9 restriction to male germ cells elicits autonomous double-strand-break repair, consequently creating offspring with diverse mutations. Thus, from a single population cross, we can obtain a deletion matrix covering a large expanse of DNA at both coarse and fine resolution. We demonstrate the ease and power of CAMIO by mapping 5’UTR sequences crucial for *chinmo’s* post-transcriptional regulation.

## Introduction

It never ceases to amaze that genomes, sequences composed of a seemingly simple four-letter code, serve as the fundamental blueprints for life. From these four nucleotides emanate layers upon layers of complexity. The central dogma depicts the basic information flow from DNA to RNA to protein. However, numerous mechanisms increase the complexity at each step and can feed backwards. Some examples include epigenetic modifications, promoter/enhancer usage, alternative splicing, post-transcriptional and post-translational modifications. Understanding how complex biology unfolds, advances, and evolves from genomic sequences requires development of precise and efficient tools to interrogate the genome. While spontaneous or induced mutations once served as the basis of our understanding about many biological processes, we now exploit sophisticated methods to directly manipulate genomes for systematic mechanistic studies.

Pioneering geneticists, like Dr. Barbara McClintock, enlightened us with fundamental understanding of genetic information and regulation at the level of genes and chromosomes. More recently, to understand gene regulation, molecular biologists have relied heavily on *in vitro* or cell culture-based assays to assess the function of DNA fragments. One example of this is promoter bashing used to pinpoint critical or regulatory sequence elements in a promoter^1, 2^. Also, in enhancer screening, larger DNA fragments upstream or downstream of a target gene are fused to reporter genes^3–5^. This synthetic approach often fails to recapitulate full endogenous patterns, and thus requires second-round rescue experiments to draw solid conclusions. Such indirect methods are inefficient and non-physiological, prompting the search for tailor-made sequence-specific DNA nucleases that permit targeted mutagenesis of native sequences.

In the early 2000s, zinc-finger nucleases (ZFNs)^6^, the first of the tailored nucleases, were heavily adopted^7, 8^. Although it seemed a promising strategy for genome editing, major setbacks, especially off-target toxicity limited its utility^9^. Therefore, the enthusiasm for ZFNs declined after the arrival of transcription activator-like effector nucleases (TALENs)^10–12^. The higher specificity of TALENs led to fewer off-target disruptions, and hence less cytotoxicity^13^. However, constructing a TALEN is technically challenging because the homologous sequences encoding the TALEN repeats are prone to recombine with each other^9^.

The newest technology, CRISPR (clustered regularly interspaced short palindromic repeats), was first exploited for targeted mutagenesis^14^ and then swiftly adopted to edit genomes of diverse organisms^15–20^. CRISPR’s popularity lies in its simplicity: a Cas9 nuclease plus an easily made guide RNA (gRNA) induces a double-strand break (DSB) targeted by DNA-RNA base pairing^14, 16^. Cells repair the DSBs either through nonhomologous end joining (NHEJ)^21^, leading to insertion or deletion (indel), or through homology-directed repair (HDR)^22^ which replaces discontinuous sequences with an available template. The fact that CRISPR can produce targeted DNA modifications with near unlimited specificity has ensured its rapid expansion throughout the biological and biomedical fields^23^. Indeed, CRISPR technology has already transformed studies from stem cell research and cancer biology to food production and pest control^24^.

*Drosophila melanogaster* has been a powerful model organism for decoding the genome to understand complex biology^25^. However, even with existing genetic tools, it remains quite challenging to interrogate the entire fly genome, especially non-coding regions. Enhancing fly genetics with CRISPR is particularly needed for large-scale genome-wide screens as well as focused, detailed sequence analyses^26^.

Here we describe a new high-throughput technology for deletion analysis called CAMIO (<u>C</u>as9-mediated <u>A</u>rrayed <u>M</u>utagenesis of Individual <u>O</u>ffspring). We built a CRISPR-based mutagenesis pipeline in *Drosophila* male germ cells, to achieve massive production of independent indels in targeted loci with germline transmission^27, 28^. We further created a transgenic system for simultaneously targeting multiple sites with an array of gRNAs. This way, we can readily generate a huge collection of organisms harboring either diverse, small, localized indels or large, defined deletions (inter-target deletions). This will enable efficient deletion analysis of sizable genomic regions *in vivo*. We include here an example that demonstrates the power of CAMIO. In 2006 our lab discovered that the expression of temporal protein Chinmo was regulated via its 5’UTR^29^. Using CAMIO, we are able to rapidly ascribe a critical aspect of this temporal control to a 300-bp sequence of the 2kb UTR. In conclusion, CAMIO enables high-throughput organism-level genome structure-functional studies.

## Results

### Independent targeted mutagenesis in individual male germ cells

To mutate the genome in a high-throughput manner requires a germline pipeline for independent mutagenesis in individual germ cells rather than germline stem cells (GSCs). We have shown that the *bam* promoter can effectively and specifically drive flippase induction in female germ cells, and not in GSCs^28^. *bam*’s restriction to germ cells in both the male and female germline prompted us to explore whether we could conduct independent mutagenesis in individual germ cells.

To this end, we made *bamP(898)-Cas9* and tested its ability to mutate gRNA-targeted sites in male as well as female germline. For proof of principle, we chose the *ebony* gene for targeted mutagenesis. Loss-of-*ebony* mutations are easy to detect and there was a readily available transgenic gRNA targeting *ebony* (Fig. 1a)^19^. We determined the mutagenesis efficiency in male versus female founders (Fig. 1b). Surprisingly, over 25% of male gametes as opposed to only ∼3% of female gametes carried loss-of-function *ebony* mutations. This result demonstrates that the male germline is particularly susceptible to *bamP-Cas9*-mediated genome editing.

**Figure 1.**
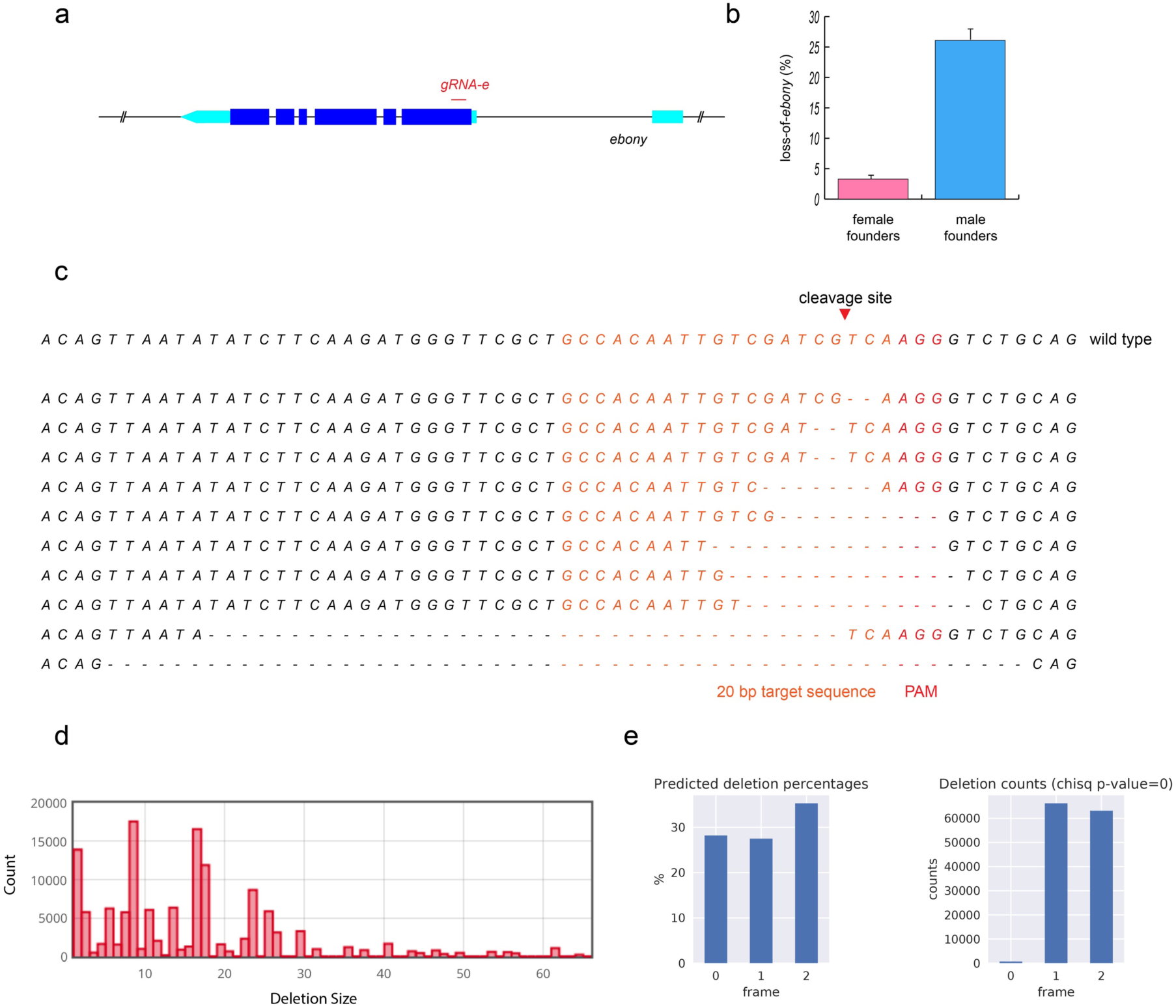
*bamP-Cas9* induces efficient CRISPR targeted mutagenesis in male germ cells. (A) Ebony transcript shown with UTRs in turquoise. U6 drives guide RNA (*gRNA-e*), which targets 5’ end of *ebony* coding sequence. (B) Percent of progeny with ebony loss-of-function (LOF) mutations from female or male founders. Mean ± SEM (n=9). (C) Sequences of 10 randomly selected *ebony* LOF progeny from the same male founder. (D) Deletion profile of *UAS-shibire* transgene following CRISPR targeting collected from 919 phenotypically wildtype progeny. NGS data was analyzed and presented by Cas-analyzer, www.rgenome.net^30^. (E) Left, percentage of predicted indels (calculated by FORECasT^31^ using *gRNA-shi* and *UAS-shibire* sequences) grouped into 3 reading frames, 0 represents the in-frame indels. Right) Actual reading frame percentages from phenotypically wildtype progeny, calculated from mapping results (obtained with CAS-analyzer). Chi-square test assuming equal distribution was used to assess the significance. The result is below machine precision and thus set to zero.

To address if the Cas9-mediated editing events occurred independently, we sequenced a part of the *ebony* locus in individual mutants carrying *ebony* deficiency. From a single male founder, we analyzed ten progeny with *ebony* loss-of-function phenotypes. We uncovered seven different indels around the Cas9 cut site (Fig. 1c). One identical *ebony* mutation occurred in three siblings, possibly resulting from either differential deletion of GTC repeats or from microhomology-mediated repair^32^. The recovery of many distinct indels from a single founder argues for independent Cas9 actions in individual germ cells. This result encouraged us to establish *bamP* induced CRISPR in the male germline as a ‘targeted mutagenesis pipeline’.

A mutagenesis pipeline could be useful for producing novel alleles of a protein of interest. To explore this idea, we tested CRISPR’s usefulness to produce novel alleles of a well characterized gene, *shibire. shibire* encodes *Drosophila* Dynamin, a motor protein crucial for synaptic vesicle endocytosis^33^. *shi^ts1^*, a temperature sensitive allele containing a missense mutation, is widely used in behavioral assays to temporarily shut off neuronal activity^34^. We therefore designed a gRNA against the *UAS-shibire* transgene at the region where the temperature-sensitive *shi^ts1^* point mutation is located, presuming that we could produce additional temperature-sensitive or dominant negative alleles.

We tested our mutagenized transgene by expressing it in the eye with GMR-Gal4 and screening at 29 degrees. Unfortunately, the rough eye phenotype was not confined to temperature-sensitive or dominant negative alleles, but was also the result of high transgene expression in the eye. Frameshift mutations in the beginning of *shibire* would lead to premature stop codons, and the resultant small truncated proteins are likely non-functional. By contrast, in-frame mutations would create essentially full-size proteins. However, loss of critical amino acids could disrupt key catalytic functions but preserve the protein’s ability to polymerize, thus creating a dominant negative allele. We therefore surveyed phenotypically wildtype offspring to see if they lacked in-frame mutations. We pooled around 1000 phenotypically normal progeny collected from 20 male founders for amplicon analysis with next generation sequencing (NGS). We obtained a large collection of diverse indels, with the majority of deletions smaller than 30 bps (Fig. 1d). Notably, there is a clear under-representation of in-frame mutations (Fig. 1e). The selective loss of such in-frame mutations is noteworthy, and supports the feasibility of making ‘novel’ useful proteins via deleting various amino acids of interest in a high-throughput manner.

Taken together, our data demonstrate that *bamP(898)* effectively restricts Cas9-mediated mutagenesis to germ cells. There is no evidence that clonal expansion contributes to the exceptionally high mutation efficiency in male founders.

Therefore, transgenic CRISPR, induced by *bamP*, can effectively serve as a pipeline for mass production of targeted mutations.

### CAMIO: Cas9-mediated arrayed mutagenesis of individual offspring

Despite independent mutagenesis in each germ cell, using only one gRNA limits the offspring variation, as all indels are anchored around the same Cas9 cut site. To expand the diversity of deletions that one can recover from a single population cross, we next explored the possibility of multiplexing gRNA-targeted mutagenesis. Our vision for multiplexing gRNAs is to have a collection of gRNAs from which one is stochastically selected, rather than simultaneously expressing multiple gRNAs. Incorporating this multiplexed design into the male germline in combination with *bamP(898)*-Cas9 would enable both stochastically chosen gRNAs and offspring independent mutations. Supplying one gRNA at a time prevents contamination of rare deletions by much more frequent second-site mutations. This way, discrete clusters of simple deletions can be recovered from a single population cross. Therefore, we can tile a sizable DNA region with diverse small deletions with a repertoire of evenly spaced gRNAs.

To examine the feasibility of our multiplexing design, we targeted a *UAS-mCD8::GFP* transgene with four independent gRNAs. To stochastically activate only one out of the four gRNAs, we made a conditional U6-gRNA(x4) transgene that is dependent on PhiC31-mediated recombination (Fig 2a). Using the *nos* promoter, we control the induction of the phiC31 recombinase in GSCs. Thus, in each of the 12-24 GSCs per male fly founder^35^, the transgene is irreversibly recombined to express a single gRNA. Recombination occurs between a single attB site downstream of the U6 promoter and a choice of attP sites upstream of each gRNA; once reconstituted, the ubiquitous U6 promoter drives expression of only one of the gRNAs. The intra-chromosomal recombination excises an intervening 3xP3-RFP. Given the rather small size of each gRNA as compared to the large 3xP3-RFP, the differences in length between the attB site and the choice of any one attP site is therefore relatively trivial. Based on a previous, similar construct for multicolor imaging^36^, we expect that each gRNA should be expressed at comparable frequencies. For brevity, we name the conditional U6-gRNA transgene pCis, and then, in braces, add the number of gRNAs and the name of the targeted DNA. For example, to target the UAS-mCD8::GFP transgene, we created pCis-{4gRNAs_mCD8}. Also, when we describe the individual gRNAs, we number them in sequence from 5’ to 3’.

**Figure 2.**
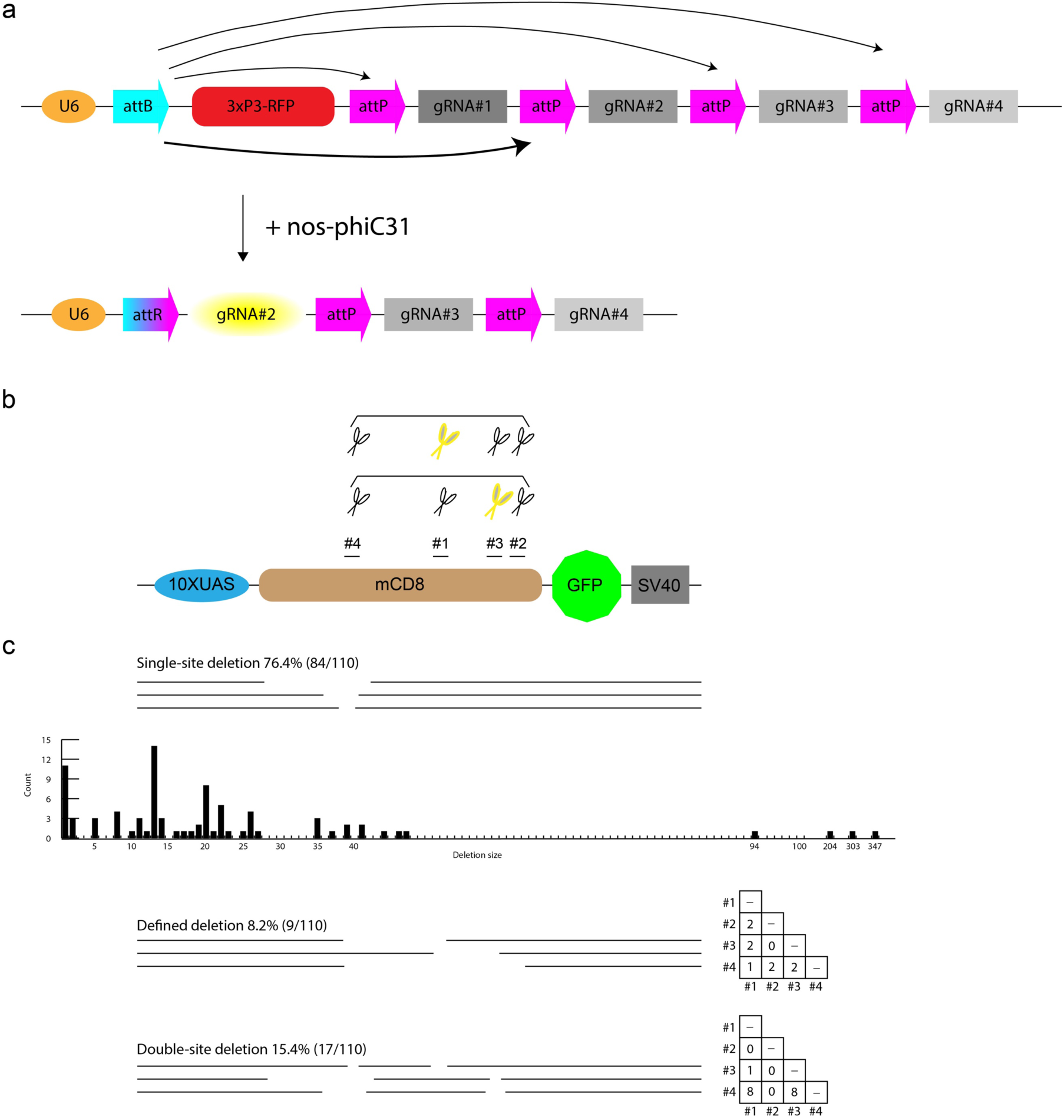
CAMIO produces diverse arrayed mutations around selected gRNA target sites. (A) Schematic of a conditional U6 gRNA transgene: pCis-{4gRNAs_mCD8}. The U6 promoter is separated from the gRNAs by a large fragment containing a 3xP3-RFP marker. With phiC31 recombination, the attachment site, attB can recombine stochastically with any of the four attP sites (black arrows), creating attR. The phiC31 recombinase is expressed in GSCs, controlled by *nos.* After recombination, a single gRNA is under the control of the U6 promoter (yellow oval) (B) 4 gRNA target sites along the mCD8 coding sequence were selected for pCis-{4gRNAs_mCD8} to disrupt mCD8::GFP expression. Simple or combined multiplexing mutagenesis can be achieved by incorporating one or more pCis-{4gRNAs_mCD8} transgenes. Here, we show two transgenes, each represented by four pair of scissors. Stochastically chosen gRNAs in each set are further marked in yellow. (C) Three categories of deletions arose from applying two copies of pCis-{4gRNAs_mCD8}: single-site deletion, defined deletion, and double-site deletion. Deletion size and count are depicted for single-site deletions. Right, use matrix of gRNA choices, deduced from sequencing GFP negative progeny.

For the multiplexed targeted mutagenesis of mCD8::GFP, we established male founders carrying *UAS-mCD8::GFP*, *bamP(898)-Cas9*, *nos-phiC31*, and *pCis-{4gRNAs_mCD8}*, and crossed them to *act5C-Gal4* females for easy scoring of GFP fluorescence in the progeny. Overall, approximately 35% of the progeny lost GFP expression. We collected 30 GFP-negative offspring from two founder males. Sequencing the mCD8-coding region revealed that each GFP-negative offspring carried an indel corresponding to a single gRNA (Supplementary Fig. 1). Encouragingly, we recovered various deletions resulting from activation of each of the four gRNAs. However, the frequency of mutations at each target site varied. Both founders yielded many more deletions around the gRNA#1/#4 targets than the gRNA#2/#3 targets, possibly reflecting their different on-target potencies.

We found that the majority (83.3%) of deletions removed 20 or fewer bp and that the largest one eliminated 85 bp. To tile a sizable DNA region with such small deletions would require many gRNAs bombarding the region of interest at a density of around one gRNA per 100 bp. Alternatively, we should be able to create larger deletions spanning two Cas9 cuts elicited by two gRNAs acting at a distance. To explore co-employment of two gRNAs, we provided two copies of *pCis-{4gRNAs_mCD8}* for multiplexed dual mutagenesis of *UAS-mCD8::GFP* (Fig. 2b). We obtained a comparable loss-of-GFP mutation rate at ∼35% despite co-expressing two identical or distinct U6-gRNAs. This phenomenon implicates that Cas9 activity (either level or duration) limits the efficiency of gRNA-directed mutagenesis in germ cells. Nonetheless, we could recover various diverse mutations from the dual gRNA-derived GFP-negative progeny, including many single-site deletions (76.4%) and quite a few double-site deletions (two target sites with independent indels; 15.4%) as well as some large deletions spanning two gRNA target sites (inter-target deletions; 8.2%) (Fig. 2c). Notably, the single-site deletions greatly outnumbered those involving two sites. This outcome is favored in large-scale deletion analysis, as it increases the chance of recovering deletions without second-site contamination.

The above results demonstrate that using dual gRNA sets enables us to tile a region of interest not only with indels, but also with defined deletions. Random selection of a single gRNA from each of the two identical sets which contain four gRNAs will yield six possible defined deletions. Encouragingly, from a collection of only nine inter-target deletions, we recovered five of the six anticipated defined deletions. Nevertheless, physical hindrance may prevent two Cas9 complexes from acting simultaneously on very close gRNA targets. This may explain why we failed to recover the smallest defined deletion of 37 bp between the Cas9 cut sites of gRNA#2 and #3 targets. These results suggest that inter-target deletions utilizing two gRNAs can support rapid systematic DNA deletion analysis.

In sum, we established an effective strategy to express various permutations of two gRNAs in male GSCs. In combination with restricting Cas9 to male germ cells, we built a germline pipeline for multiplex targeted mutagenesis. We dub this genetic system CAMIO (Cas9-mediated arrayed mutagenesis of individual offspring), which can derive from a single population cross a matrix of variable deletions. This strategy enables deletion analysis of a substantial DNA region with both coarse (inter-target deletions) and fine (a variety of single- or double-site deletions) resolution. Below, we prove the power of CAMIO in structure-functional analysis of a 2.2kb-long genomic fragment.

### Structure-functional analysis of *chinmo* 5’UTR

The Chinmo BTB-zinc finger nuclear protein is dynamically expressed in intricate spatiotemporal patterns in the developing *Drosophila* central nervous system. Such dynamic Chinmo expression governs various aspects of temporally patterned neurogenesis, including age-dependent neural stem cell proliferation^37, 38^ and birth order-dependent neuronal cell fate^29, 39, 40^. Notably, *chinmo* transcripts exist much more broadly than Chinmo proteins, indicating involvement of negative post-transcriptional regulation^29, 38^. Consistent with this notion, *chinmo* transcripts have long UTRs, including a 2.2kb 5’UTR and an 8.5kb 3’UTR^41^. To locate the involved regulatory elements in such long UTRs by conventional structure-functional analysis would be a daunting task.

Nonetheless, we started by making reporter transgenes carrying *chinmo* 5’ and/or 3’ UTR(s). For a functional readout, we utilized the development of the *Drosophila* mushroom body (MB), which involves an orderly production of γ, α’β’, and αβ neurons. We first determined the roles of the 5’ vs. 3’UTR in downregulation of Chinmo expression along MB neurogenesis^29^, by examining the change in expression of reporter transgenes (containing either or both UTRs) from early to late larval stages (Supplementary Fig. 2). Notably, presence of the *chinmo* 5’UTR drastically suppressed the reporter expression. Interestingly, only in the absence of the 3’UTR did we detect an enhanced 5’UTR-dependent suppression at the late larval stage. These phenomena ascribe the *chinmo* downregulatory mechanism(s) to the 5’UTR, and unexpectedly revealed some upregulation by the 3’UTR. This upregulation could potentially be a transgene-specific artifact, arguing for the importance of performing assessments in the native environment. We thus turned to CAMIO to carry out structure-functional analysis of the native *chinmo* 5’UTR.

*chinmo*’s 5’UTR is separated into 3 exons; the first two exons are neighboring and the distant 3^rd^ exon is separated from the 2^nd^ by 36kb (Fig. 3a). A gRNA set was designed to target each exon for CRISPR mutagenesis (Fig. 3b). We provided two copies of the same set for induction of both indels and inter-target deletions within an exon. Additionally, we paired gRNA sets for exon 1 and 2 to create larger deletions that span the exon1/2 junction. Thus, we could delete various parts of *chinmo* 5’UTR in its endogenous locus with simple fly pushing.

**Figure 3.**
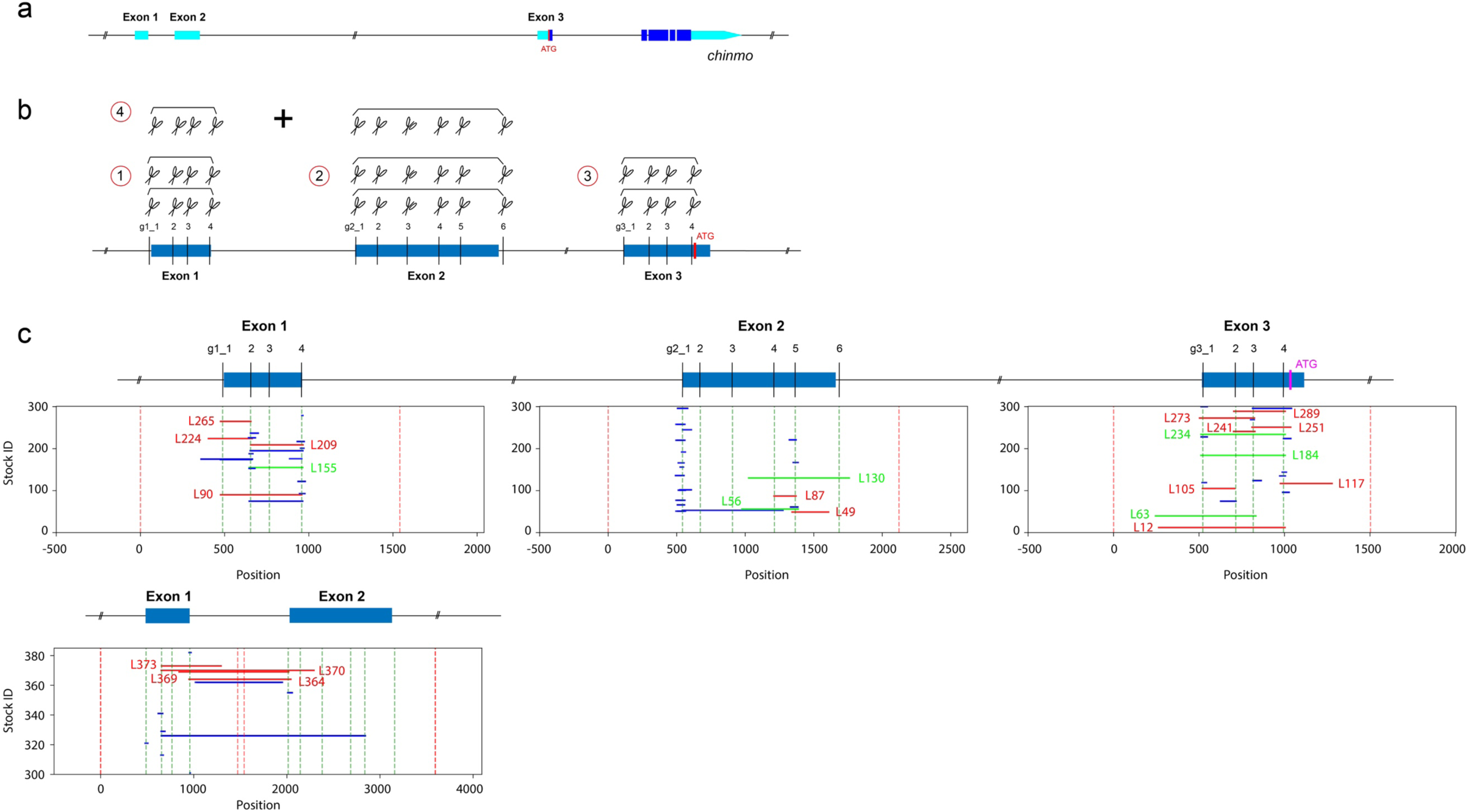
Applying CAMIO on *chinmo* 5’ UTR. (A) An illustration of the *chinmo* gene, UTRs are depicted in turquoise. (B) We selected 4 target sites on *chinmo* exon 1 (455 bps), 6 target sites on exon 2 (1061 bps), and 4 target sites on exon 3 (639 bps). Each exon was dissected by CAMIO with a pair of gRNA sets, depicted as a set of scissors. Additionally, we combined gRNA sets from exon1 and exon2. (C) Representation of larger deletions predicted by NGS of 984 offspring (300 each for individual exons and 84 for Ex1-Ex2). Predicted deletions over 100 bps were subject to Sanger sequencing for confirmation. Confirmed >100 bps deletions are marked in red and given an ID#. Selected homozygous viable deletions (marked in green) that cover larger proportions of each exons were selected for MB development studies.

Based on the previously observed deletion rate of around 35%, we collected 300 male progeny from each CAMIO genetic cross. We hoped to saturate each 5’UTR exon with ∼100 different deletions. In total, 1200 CAMIO males were harvested from the four different gRNA array combinations (exons 1, 2, 3, and 1+2). We mapped potential indels by sequencing indexed PCR products in a high-throughput manner.

We detected numerous indels around each gRNA target site (Supplementary Fig. 3) and also recovered many inter-target deletions that together allow efficient coverage of the entire 5’UTR (Fig. 3c). We made organisms homozygous for the large inter-target deletions and examined MB morphology. Markedly, we found similar aberrant MB morphology with two exon 2 inter-target deletions, *chinmo^Ex2L56^* and *chinmo^Ex2L130^* (Fig. 4a). These deletions overlap by ∼300 bp. Variable defects in the perpendicular projection of the bifurcated αβ axon lobes appeared at comparable frequencies (∼30-50%) in homozygous as well as transheterozygous brains. Further, the penetrance of this phenotype is sensitive to Chinmo dosage, as a *chinmo* deficiency line effectively suppressed the phenotype (Fig. 4b). Marchetti and Tavosanis recently proposed that Chinmo downregulation plays a role in promoting α’β’ to αβ MB neuron fate transition at the prepupal stage^42^. Therefore, we assessed Chinmo levels, and observed aberrantly elevated Chinmo in young MB neurons around pupa formation in both *chinmo^Ex2L56^* and *chinmo^Ex2L130^* homozygous mutants (Fig. 4c). This is consistent with the notion that these overlapping deletions have uncovered the essential region for this prepupal downregulation of Chinmo. In summary, a single round of CAMIO allowed us to identify a 300 bp locus in the 2.2kb 5’UTR critical for proper MB development.

**Figure 4.**
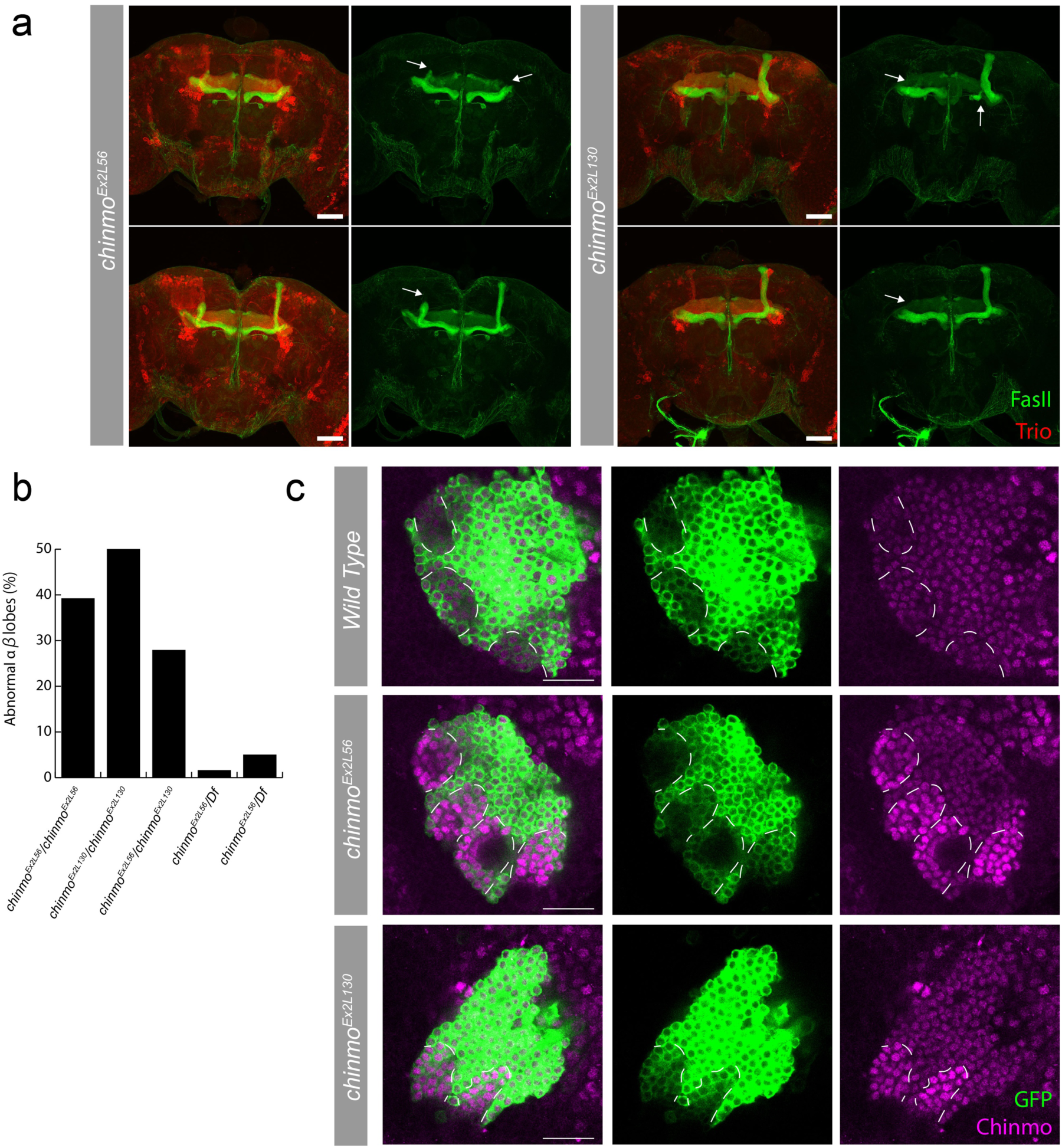
Overlapping deletions in 5’UTR alter Chinmo expression and MB morphology. (A) Stacked confocal images of adult MBs stained for FasII (green, αβ and γ lobes) and Trio (red, α’β’ and γ lobes). Missing or misshapen αβ lobes (arrows). Scale bars: 50 μm. (B) Percent of flies with abnormal αβ lobes. Homozygous and transheterozygous deletions have strong MB αβ lobe defects, whereas hemizygous deletions over a *chinmo* deficiency line (*Df*) are much less penetrant. (C) Single optical sections of MB lineages (*OK107-Gal4*, *UAS-mCD8::GFP*) immunostained for GFP and Chinmo at white pupa stage. Chinmo staining is elevated in newly derived MB neurons (weaker GFP signal, outlined by white dashed lines) in homozygous *chinmo^Ex2L56^* and *chinmo^Ex2L130^*. Green: GFP; Magenta: Chinmo. Scale bars: 20 μm.

Meanwhile, notably absent were indels or large deletions involving the target sites g2_2 and g2_3. This area in exon 2 may carry essential sequences for Chinmo regulation that is critical for organism viability. Alternative explanations for the failure in recovering indels from that region include: a relatively shallow sequencing depth of the exon 2 region, our small sample size, unexpectedly low gRNA on-target strength for these gRNAs, or flawed design in the exon 2 gRNA set pCis-{6gRNAs_chinmo Exon2}. To address the last concern that bothered us most, we assessed the usage of specific gRNAs for the exon 2 set in progeny that do not contain Cas9 (Supplementary Fig. 4). While g2_2 and g2_3 were not recruited as frequently as others, they each still emerged 6-7% of the time. A rate of 6-7% should be sufficient for us to recover some indels, as gRNA#1 was selected ∼15% of the time and produced multiple indels in our CAMIO experiment.

To examine whether this region of the 5’UTR is indeed critical, we exploited mosaic analysis to create somatic mutations in different tissues. A transgene, *dU6_g2+3*, was hence assembled to ubiquitously express both g2_2 and g2_3. We began with MB-specific CRISPR mutagenesis by inducing *UAS-Cas9* specifically in the MB lineage using a MB specific Gal4 (*OK107-Gal4*). We saw no temporal fate changes in the MB, the classic phenotype of *chinmo* misregulation in the MB. We next elicited CRISPR mutagenesis in all neural stem cells (neuroblast: NB) with NB-restricted Cas9 (*dpn-Cas9*) and *dU6_g2+3*. These animals were viable and showed no abnormal tumor-like NBs in larval or adult brains (typical of Chinmo overexpression)^38^. These data failed to provide evidence in support of presence of critical brain regulatory elements in the region targeted by g2_2 and g2_3. We strengthened this negative conclusion by directly removing various small-to-large fragments around g2_2 and g2_3 targets from the above *chinmo* UTR-containing GFP transgene. In developing MBs, we observed indistinguishable GFP expression profiles between wild-type and modified 5’UTRs (Supplementary Fig. 5). In contrast to our negative findings in the brain, we found severe embryonic or early larval lethality when we induced early ubiquitous somatic mutations with *act5C-Cas9* and *dU6_g2+3*. This dominant lethality provides a direct explanation for our failure in recovering viable organisms carrying g2_2 or g2_3 induced indels. Together, these data suggest an essential role for *chinmo* 5’UTR outside of the brain.

In conclusion, a single round of CAMIO successfully led us to uncover two critical regions of the *chinmo* 5’ UTR. The first critical region lies around the g2_2 and g2_3 targets and carries essential sequences for organism viability, unrelated to Chinmo’s functions in the brain. The second critical region of the chinmo 5’UTR was uncovered by the overlapping *chinmo^Ex2L56^* and *chinmo^Ex2L130^* deletions. We determined that this ∼300bp region is essential to down-regulate Chinmo expression, ensuring proper MB development. This fruitful case-study exemplifies the power of CAMIO in high-throughput unbiased deletion analysis of the genome.

## Discussion

Two innovations synergistically enable CAMIO, a germline pipeline for arrayed CRISPR mutagenesis. First, the *bamP* promoter can specifically limit Cas9 endonuclease activity to individual male germ cells—thus individual offspring receive independent mutations. Second, the random-choice gRNA arrays provide extensive coverage for deletion analysis, with both small indels and large deletions. Hence, the combination of bamP-Cas9 and gRNA arrays used in CAMIO enables *in vivo* targeted deletion analysis with both minimal molecular biology and minimal fly pushing.

We were happily surprised to discover a much higher CRISPR mutagenesis rate in male compared to female germ cells using *bamP*. This sex difference was also observed in CRISPR-induced gene targeting in our effort to improve Golic+ (manuscript in preparation). Currently, we do not know what leads to this phenomenon. *bamP* has a striking similar expression pattern in both the female and male germline: absence in GSCs and an early onset of expression during the four incomplete mitoses that produce the 16-cell germline cysts^43^. One possibility is that the *bamP* activity is higher in the male than the female germline. Another possibility has to do with the sex differences in meiotic recombination— meiotic recombination does not occur in male *Drosophila*. Perhaps reduced access to homologous chromosomes as templates for homology-mediated repair favors indels. In any case, this feature allows us to utilize *bamP* to build a high-efficiency pipeline for targeted CRISPR mutagenesis in male germ cells.

Conventional gRNA multiplexing provides all gRNAs at once as a cocktail, which expands indel diversity but inevitably creates complex and often biased deletion patterns. The off-target effects of a gRNA cocktail also accumulate in an additive manner. By contrast, CAMIO selects a single gRNA from each set and complexity can be added by increasing the number of sets. Thus, CAMIO confers every gRNA with some autonomy while achieving multiplexed mutagenesis as a whole. Off-target concerns in CAMIO can be adequately addressed by examining multiple independent mutations of similar kinds. Also, arrays of targeted mutations can be introduced into specific genetic backgrounds with CAMIO. For example, when performing CAMIO on the *chinmo* 5’ UTR, we purposefully targeted a 2L chromosome arm that also carries transgenes for twin-spot MARCM ^44^. Hence, all the CAMIO *chinmo* indels were immediately ready for mosaic analysis. Independent indels can be directly screened for visible phenotypes in mosaic organisms. However, we favor mapping the indels first by NGS, which can be conducted in a high-throughput manner via combinatorial sample indexing. We have also reduced the costs by pooling distinct amplicons for co-indexing.

In general, we recovered similar indel spectrums to what has been commonly described. For gRNAs that are inherently potent, like g1_4 for *chinmo* 5’ UTR, we obtained many indels around the cut site. Despite recovering numerous single-site deletions, we rarely see single-site deletions exceeding 30 bps in length. Therefore, the observed ease in creating diverse inter-target deletions by CAMIO is particularly valuable for systematic DNA dissection. In the case of CAMIO on *chinmo* 5’UTR, we recovered most of the predicted inter-target deletions with the exception of the ‘toxic’ g2_2 and g2_3. Notably, the largest inter-target deletion we have identified exceeds 2 kb in length. These observations suggest that we can be more aggressive in choosing more disperse gRNA targets to cover larger genomic regions.

After all, the capacity of CAMIO is mainly determined by how many gRNAs one can pack into a single set. Given the small size of gRNAs, we expect no problem in packing six or even more gRNAs without compromising the system. This intuition was largely supported by seeing reasonable recruitment frequencies for all six tandem gRNAs in the *chinmo* exon 2 set. Further, there is still room for improvement on the gRNA selection process. For instance, increasing the distance between the U6 promoter and the gRNA set would make the selection more impartial. In sum, we have shown that the CAMIO system holds great promise for *in vivo* deletion analysis. Yet, our demonstrations have not nearly reached the limitations of CAMIO as far as the number of targets and size of DNA that can be evaluated in a single experiment.

We used CAMIO to perform deletion analysis on the 5’UTR of *chinmo,* which has important roles in governing Chinmo protein levels. There evidently exist multiple mechanisms governing *chinmo* expression throughout development. We successfully identified a region responsible for Chinmo downregulation in the MB around pupa formation. The resulting elevated Chinmo expression affected MB morphogenesis, possibly due to abnormal neuronal fate transition. In addition, we found a large region critical for embryo viability. While roles for Chinmo have been described in the brain and as a downstream target of JAK/STAT in the testes^45^, our data suggest another essential role for Chinmo in embryonic development. The identification of discrete non-coding regions regulating different biological processes within a single UTR has exemplified the utility of CAMIO in resolving complex UTR functions. Given its multiplex and combinatorial power, CAMIO should also greatly aid the dissection of promoters, enhancers, long non-coding RNAs, DNA repeats and more.

In theory, CAMIO should work for any organism where a *bamP*-like promoter exists. Particularly, if a founder parent (possibly a father) can produce a large number of targeted mutants, CAMIO may become a desirable genetic screening platform for interrogating the genome. A mouse gene, *Gm114*, was identified as a putative ortholog of *Drosophila bam*^46^. Encouragingly, strikingly similar to *bam, Gm114* is greatly enriched in undifferentiated spermatocytes and spermatids but absent or extremely low in undifferentiated spermatogonia. Orthologs of *bam* and *Gm114* were also found in zebrafish, chicken, macaque, and others. We have pioneered CAMIO as a germline pipeline for arrayed CRISPR mutagenesis in *Drosophila*. CAMIO can expedite systematic structure-functional analysis of the genome across diverse model organisms.

## Methods

### Fly strains

We used the following fly strains in this work: (1) *bamP(898)-Cas9* in *attP2*; (2) *U6:3-gRNA-e*^19^; (3) *Df(3R)ED10838/TM2* (BDSC #9485); (4) *dU6-3-gRNA-shi* in *attP40*; (5) *UAS-shibire* in *attP2*; (6) *GMR-Gal4*; (7) *nos-phiC31-nls #12*; (8) *10XUAS-mCD8::GFP* in *attP40*^47^; (9) *act5C-Gal4/TM6B*; (10) *pCis-{4gRNAs_mCD8}* P element insertion line #3, #10 on II, and #9, #12 on III; (11) 13XLexAop2-5’ UTR-smGFP-OLLAS *in* VK00027, 13XLexAop2 –smGFP-cMyc-*3’ UTR* in *attP40*, and *13XLexAop2-5’ UTR-smGFP-V5-3’ UTR* in *su(Hw)attP5*; (12) *41A10-KD*; (13) *DpnEE-KO-LexAp65*; (14) *pCis-{4gRNAs_chinmo Exon1}* P element insertion line #23, #25 on II, and #4, #5 on III; *pCis-{6gRNAs_chinmo Exon2}* #7, #11 on II, and #14 on III; *pCis-{4gRNAs_chinmo Exon3}* #5 on II, and #2, #3 on III; (15) *FRT40A,UAS–mCD8::GFP,UAS–Cd2-Mir/CyO,Y*^44^; (16) *OK107-Gal4*; (17) *UAS-Cas9*^19^; (18) *dU6_g2+3* in *VK00027*; (19) *Dpn-Cas9* (unpublished reagent); (20) *act5C-Cas9*^19^; (21) *13XLexAop2-53UTR-smGFP-V5-d*, *13XLexAop2-53UTR-smGFP-V5-bD*, and *13XLexAop2-53UTR-smGFP-V5-D23* in *attP40*.

### Molecular biology

To create *bamP(898)-Cas9*, the full *bam* promoter (−898)^27^ was ordered from gBlocks, IDT, and Cas9 was also flanked by *bam* 3’ UTR. To create UAS-shibire, codon-optimized *shibire* coding sequence that carries the gRNA-shi target site was ordered from GeneArt gene synthesis, and then cloned into pJFRC28^48^. To generate dU6-3-gRNAs, we replaced 10XUAS-IVS-GFP-p10 of pJFRC28 with dU6-3 promoter and gRNA scaffold fragment from pTL2^28^. For dU6-3-gRNA-shi, GTATGGGGTATCAAGCCGAT was selected as the spacer. To create dU6_g2+3, we first generated dU6-3-g2_2 and dU6-3-g2_3 separately, and cloned dU6-3-g2_2 into the backbone of dU6-3-g2_3.

To create conditional U6-gRNA set construct, pCis-{4gRNAs_mCD8}, a U6 promoter-AttB fragment was synthetized by PCR amplification from pCFD3^19^ and cloned into pCaST-elav-VP16AD, which contained the p-Element inverted repeats (Addgene, #15308). Then, we inserted a DNA fragment (Genscript) containing 4 different gRNAs targeting the mCD8 protein tag. These gRNAs were selected based on their ON and OFF target scores (Benchling). Each of these gRNAs was preceded by an AttP site and a HammerHead ribozyme^49^. Finally, a 3Xp3-RFP-polyA(*α*-tubulin) fragment was synthetized by PCR amplification, using pure genomic DNA from a fly line in which this cassette was used as a marker (Bloomington, #54590). Then, this fragment was inserted upstream of this gRNA region. In the final construct, the AttB and AttP sites were separated by a 3.7 Kb region containing an ampicillin resistance gene, an origin of replication in bacteria and the 3Xp3-RFP-polyA marker.

pCis-{gRNAs_chinmo-Exon1/Exon2/Exon3}: following the same design described above, a DNA fragment was synthetized (Genscript), which contained 4 gRNAs (6 for Exon2) either targeting the corresponding exon or the exon-intron junction. This fragment was then inserted into pCis-{4gRNAs_mCD8}, thus removing the previous gRNAs cassette.

The *Chinmo* UTRs were amplified from Drosophila genomic DNA. smGFP^50^ fused to V5, cmyc or ollas were amplified from previously existing plasmids. Standard molecular biology techniques were used to clone the smGFP fusions containing one or two *Chinmo* UTRs into 13XLexAop2 (pJFRC15,^47^). If the construct ended with *Chinmo* 3’UTR, the SV40 signal was removed from the vector backbone. *13XLexAop2-5’ UTR-GFP-3’ UTR* was further modified to create d, bD, and D23 reporters containing various deletions in the 5’ UTR.

### Drosophila genetics

#### *ebony* and UAS-shibire mutagenesis by *bamP(898)-Cas9*

Female or male founders (*bamP(898)-Cas9/U6:3-gRNA-e*) were mated to *Df(3R)ED10838/TM6B*, and chromosomes over *Df(3R)ED10838* were scored for ebony loss-of-function phenotype. In total, 709 progeny from 5 female founders and 1574 progeny from 9 male founders were collected and phenotype was assessed. Male *UAS-shibire* mutagenesis founders were crossed with *GMR-Gal4*, and the wildtype-eyed 919 progeny were sacrificed for next generation sequencing (NGS).

#### CAMIO on UAS-mCD8::GFP

Male founders (*UAS-mCD8::GFP*, *bamP(898)-Cas9*, *nos-phiC31*, and one or two copies of *pCis-{4gRNAs_mCD8}* were mated with *act5C-Gal4* females for scoring of loss of green fluorescence in the progeny. For one copy of *pCis-{4gRNAs_mCD8}*, we screened 807 progeny from 20 founder males. 30 loss-of-GFP progeny from two founders were further subject to sequence analysis. For two copies of *pCis-{4gRNAs_mCD8}*, we screened 570 progeny from 16 founders. 110 loss-of-GFP progeny were analyzed and grouped into three deletion categories.

#### CAMIO on *chinmo* 5’ UTR

After mating with females carrying a second chromosome balancer for stock keeping, 1200 male progeny from 4 CAMIO on *chinmo* 5’ UTR gRNA set combinations were sacrificed for genomic study. We designed primer sets that produce amplicons covering exon 1, 2, 3, and exons 1+2. Males from combination 1 were intentionally numbered 1-300, and their genomic amplicons were carefully matched and mixed with counterparts from combination 2 and 3. Finally, 300 DNA mixtures plus 84 amplicons from combination 4 were tagmented (Nextera XT DNA Library Prep Kit, illumina) and barcoded (Nextera XT Index Kit v2) for NGS.

### Immunohistochemistry and confocal imaging

Fly brains at indicated larval, pupal, and adult stages were dissected, fixed, and immunostained as described previously^29, 44^. The following primary antibodies were used in this study: chicken GFP polyclonal antibody (1:500, Invitrogen, A10262); rat anti-Deadpan (1:100, abcam, ab195173); mouse 1D4 anti-Fasciclin II (1:50, DSHB); rabbit polyclonal anti-Trio (1:1000)^51^; rat anti-Chinmo (1:500, a gift from the Sokol lab)^41^. Secondary antibodies from Molecular Probes were all used in a 1:200 dilution. The immunofluorescent signals were collected using Zeiss LSM 880 confocal microscope and processed using Fiji and Adobe Illustrator.

### Bioinformatics

Sequence reads (FASTQ data) were first processed with cutadapt [https://github.com/marcelm/cutadapt] to remove adapter sequences with options: --overlap=7 --minimum-length=30 -a “CTGTCTCTTATACACATCTCTGAGCGGGCTGGCAAGGCAGACCG”. Then they were mapped to the genomic sequence corresponding to the *chinmo* region using Bowtie2^52^ with the following options: --local --score-min G,20,0 -D 20 -R 3 - L 20 -i S,1,0.50 --no-unal. Resulting SAM files were parsed with Pysam [https://github.com/pysam-developers/pysam] to extract deletion/insertion information from the Cigar strings. When the cigar string contained ‘D’ or ‘N’, we extracted mapped sequences as deletions and designated them as type D. When the cigar string contained ‘I’, we extracted them as insertions and designated them as type I. While parsing the SAM file, soft clipped reads (reads partially mapped to the genome) were detected and clipped (unmapped) portions were set aside in a FASTA file. The sequences in this FASTA file were then re-mapped using bowtie2 to the genomic sequence encompassing the *chinmo* gene. Then, mapped portions of the first mapping and that of the second mapping (when the second one existed) were merged to form a deletion event which we denoted as type L (large gap). This type of gap often contained inserted sequences in the middle. We discarded any events with less than 10 reads.

## Acknowledgements

We thank Janelia Fly Core and Quantitative Genomics for technical support. We thank Crystal Di Pietro and Kathryn Miller for administrative support. This work was supported by Howard Hughes Medical Institute.

## Author contributions

H.-M.C. and T.L. conceived the project. H.-M.C. performed the CAMIO experiments. J.G.M. conceptualized the design of CAMIO gRNA multiplexing and generated the CAMIO gRNA sets. K.S., H.-M.C., and T.L. contributed to the design, execution, and analyses of the CAMIO NGS project. D.W. contributed to the sample preparation for CAMIO NGS. R.L.M. designed the chinmo-UTRs reporter constructs. K.S. provided statistical data analyses. H.-M.C., R.L.M. and T.L. wrote the manuscript. T.L. supervised the project.

## Supplementary information

### Supplementary figure legends

**Supplementary Figure 1.**
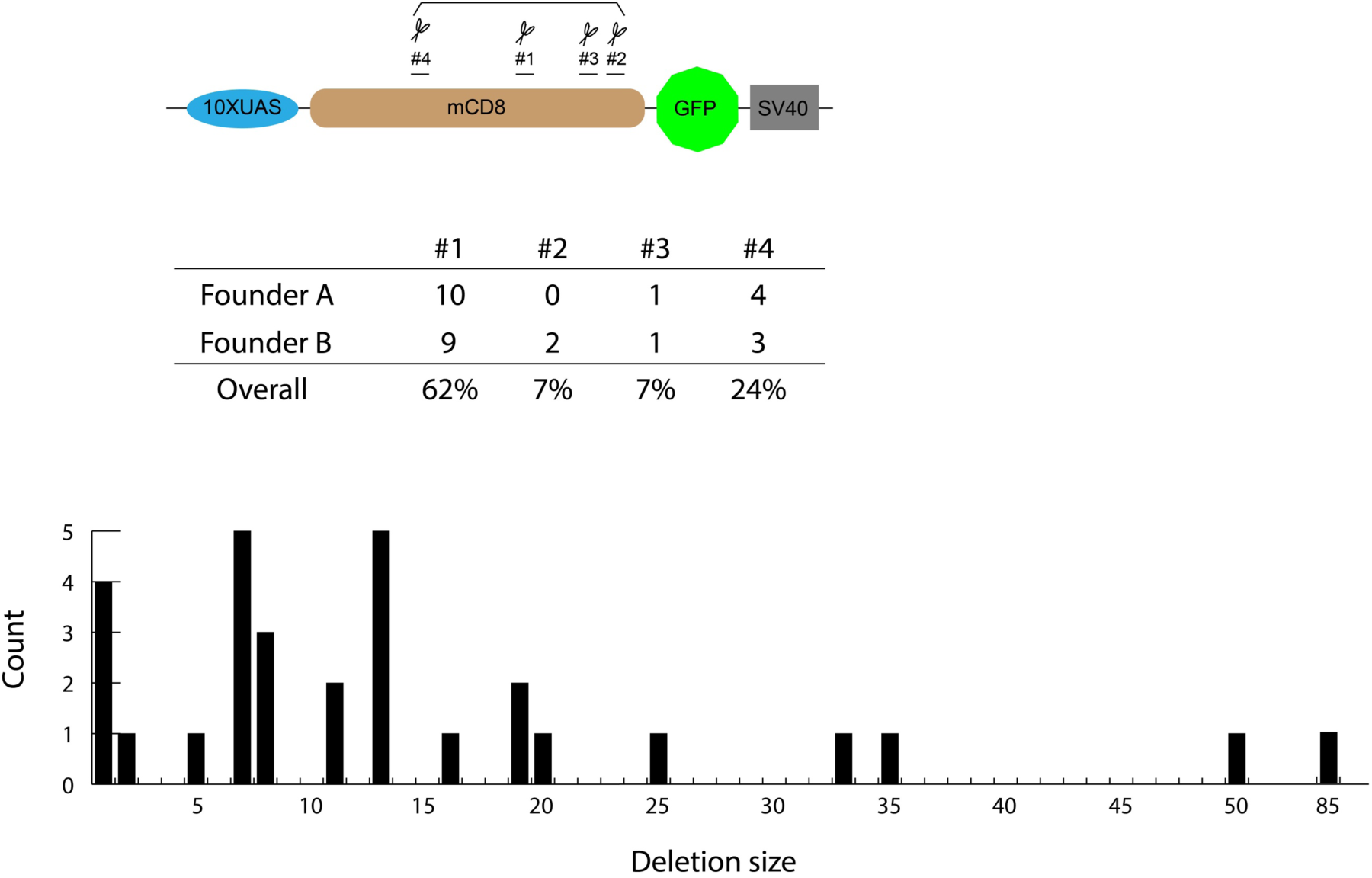
Targeted mutagenesis of mCD8::GFP with pCis-{4gRNAs_mCD8}. pCis-{4gRNAs_mCD8} was created to target four different sites along the mCD8 coding sequence of UAS-mCD8::GFP. We analyzed 30 GFP-negative progeny from two different male founders. Each GFP-negative progeny carried a indel located at one of the four target sites. The location, size, and overall counts of these 30 indels are summarized here.

**Supplementary Figure 2.**
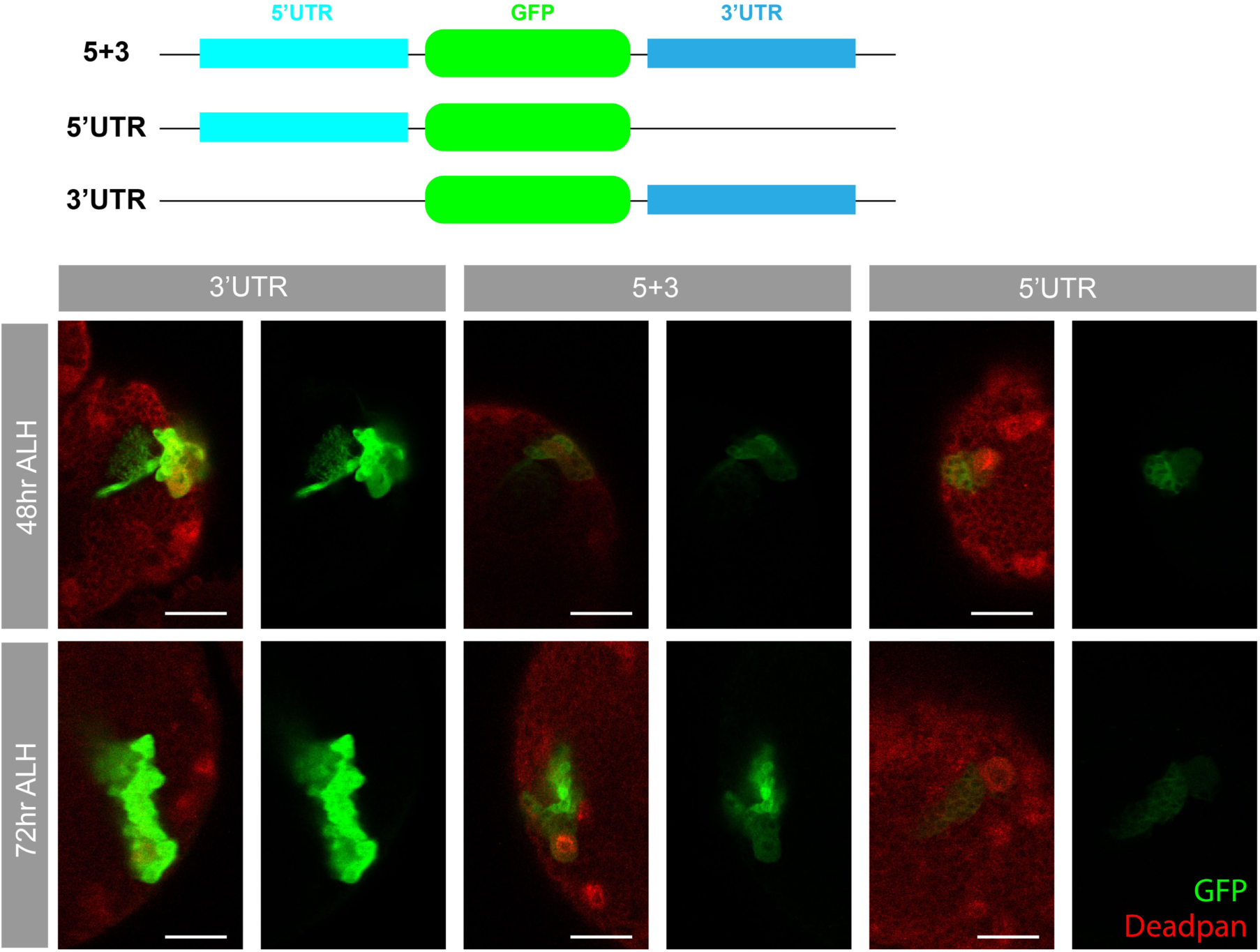
Three *13XLexAop2-GFP* reporters for investigating *chinmo* 5’ and 3’ UTRs. Top: diagram of 13XLexAop2-GFP reporters flanked by both *chinmo* 5’ and 3’ UTRs, only the 5’ UTR, or only the 3’ UTR. Bottom: GFP transgene expression was induced and restricted in MB lineages by immortalizing a transient MB NB expression (*41A10-KD*) into a sustained MB NB production of these three GFPs, with this genetic setup: *DpnEE-KO-LexAp65; 13XLexAop2-GFP-UTRs; 41A10-KD*. Their expression profile is shown by immunostaining for GFP at two time points (48 and 72 hours <u>a</u>fter larval <u>h</u>atching, ALH). GFP with the 3’ UTR is expressed at a much higher level than GFP with both 5’ and 3’ UTRs. Expression of GFP with the 5’ UTR decreases over time. Green: GFP; Red: Deadpan. Scale bars: 20 μm.

**Supplementary Figure 3.**
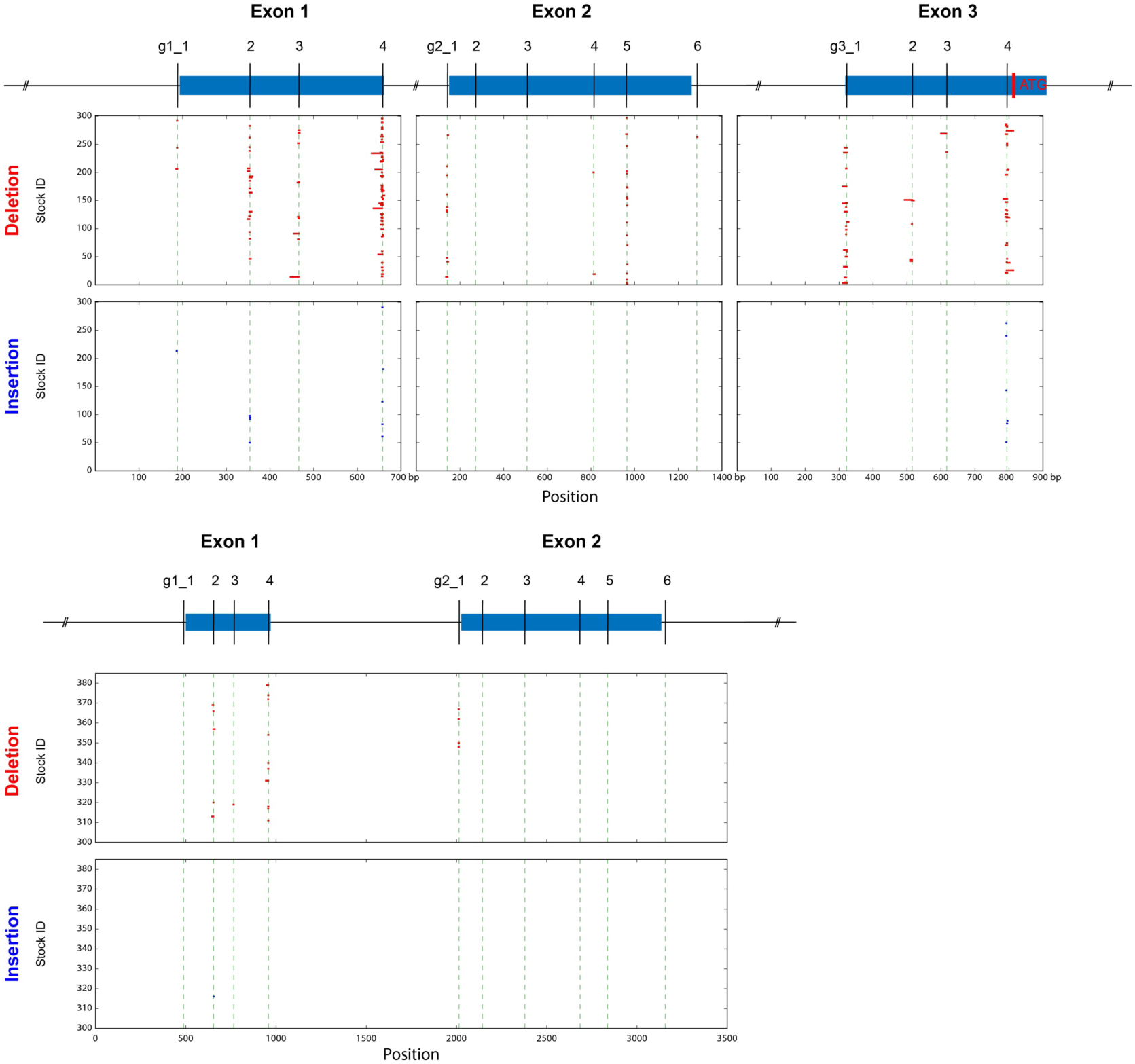
The small indels created by CAMIO in *chinmo* Exon 1, 2, and 3. We sequenced 984 progeny (300 each for individual exons, and additional 84 for Ex1-Ex2) from our CAMIO on *chinmo* 5’ UTR experiment. The size, location, and corresponding stock ID of the predicted deletions (marked in red) and insertions (marked in blue) are depicted here.

**Supplementary Figure 4.**
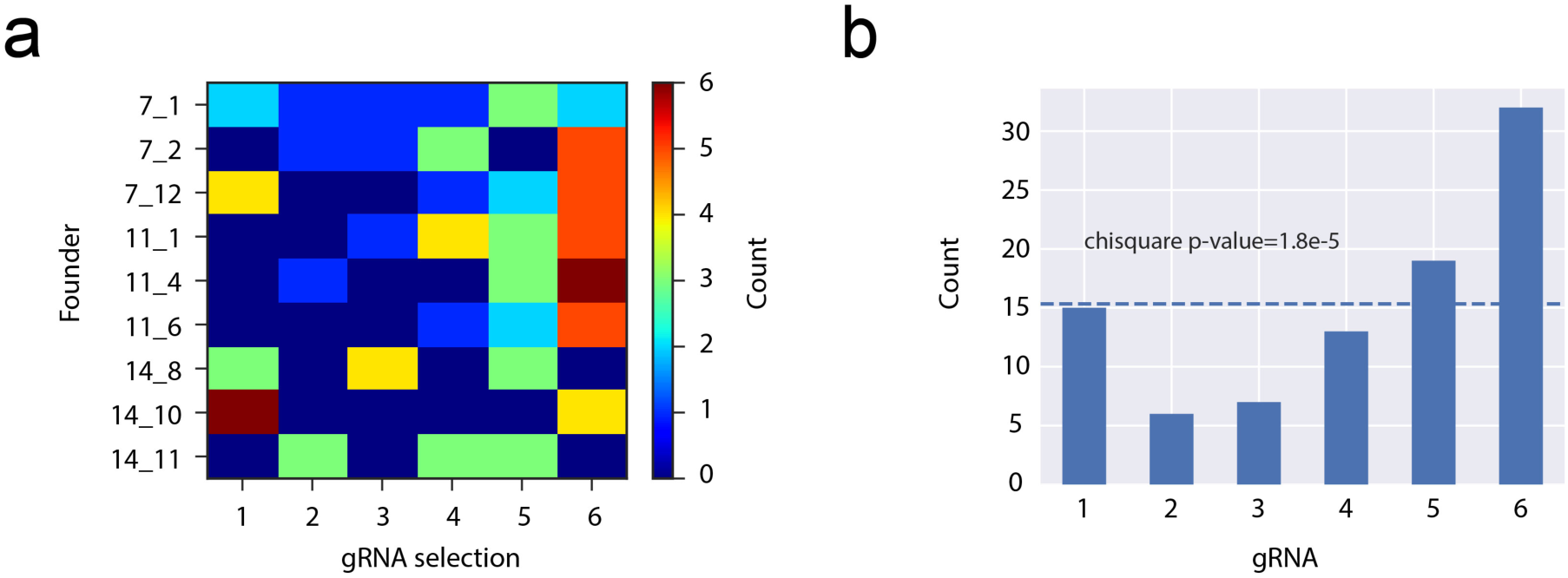
Examining gRNA array selection of *pCis-(6gRNAs_chinmo Exon2)*. We studied how frequently each gRNA gets selected after nos-phiC31 mediated recombination by scoring progeny of male *nos-phiC31; pCis-(6gRNAs_chinmo Exon2*). (A) gRNA choices of 92 progeny from 9 male founders are presented in this heat map. (B) Counts for each gRNA from all the founders are aggregated and whether there is bias in selection is tested by Chi-square test assuming equal distribution. g2_2 (6.5%, 6/92) and g2_3 (7.6%, 7/92) are underrepresented.

**Supplementary Figure 5.**
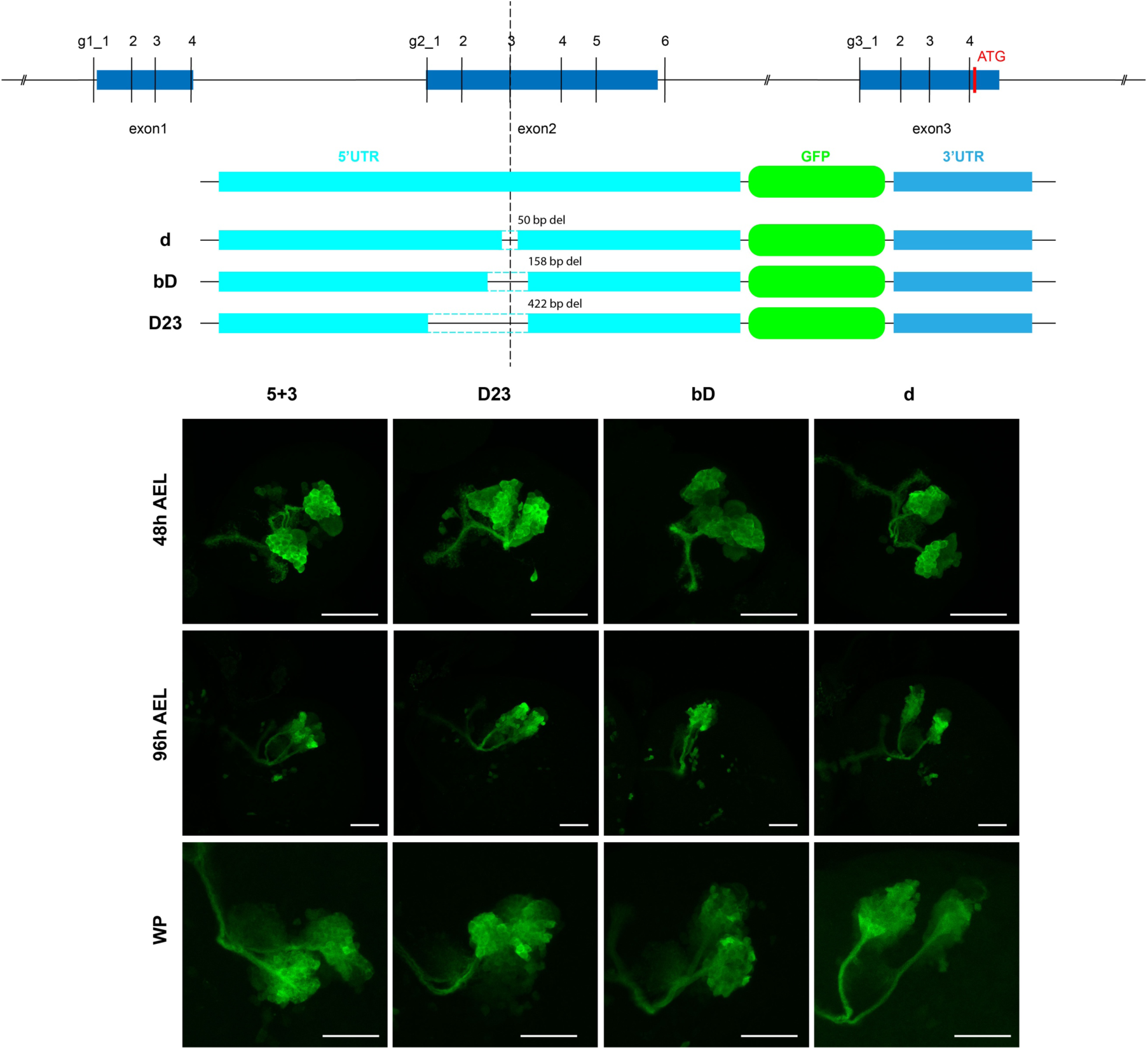
Three 5’UTR-GFP-3’UTR reporters carrying small to large *chinmo* Exon 2 deletions for studying the region around g2_2 and g2_3 target sites. We generated three additional chinmo 5’ UTR-GFP-3’ UTR reporters: d (50 bps deletion around g2_3), bD (larger 158 bps deletion), and D23 (removing all the Exon 2 sequence upstream of g2_3). GFPs were induced in the same *41A10-KD* immortalization strategy. We did not observe difference in expression among the four GFP UTR reporters at three development time points: 48h AEL (after egg laying), 96h AEL and, white pupa. Scale bars: 30 μm.

